# Trehalose 6-phosphate Controls Seed Filling by Inducing Auxin Biosynthesis

**DOI:** 10.1101/752915

**Authors:** Tobias Meitzel, Ruslana Radchuk, Erin L. McAdam, Ina Thormählen, Regina Feil, Eberhard Munz, Alexander Hilo, Peter Geigenberger, John J. Ross, John E. Lunn, Ljudmilla Borisjuk

**Affiliations:** Leibniz Institute of Plant Genetics and Crop Plant Research, Corrensstr. 3, 06466 Stadt Seeland OT Gatersleben, Germany; School of Biological Sciences, University of Tasmania, Sandy Bay, 7001, Australia; Ludwig Maximilians University of Munich, Faculty of Biology, Großhaderner Str. 2, 82152 Planegg-Martinsried, Germany; Max Planck Institute of Molecular Plant Physiology, Am Mühlenberg 1, 14476 Potsdam, Germany

**Author notes:** Correspondence: Tobias Meitzel.

**Keywords:** Trehalose 6-phosphate, auxin, sugar signaling, embryo development, seed filling, starch biosynthesis, pea

## Abstract

Plants undergo several developmental transitions during their life cycle. One of these, the differentiation of the young embryo from a meristem-like structure into a highly-specialized storage organ, is vital to the formation of a viable seed. For crops in which the seed itself is the end product, effective accumulation of storage compounds is of economic relevance, defining the quantity and nutritive value of the harvest yield. However, the regulatory networks underpinning the phase transition into seed filling are poorly understood. Here we show that trehalose 6-phosphate (T6P), which functions as a signal for sucrose availability in plants, mediates seed filling processes in seeds of the garden pea (*Pisum sativum*), a key grain legume. Seeds deficient in T6P are compromised in size and starch production, resembling the wrinkled seeds studied by Gregor Mendel. We show also that T6P exerts these effects by stimulating the biosynthesis of the pivotal plant hormone, auxin. We found that T6P promotes the expression of the auxin biosynthesis gene *TRYPTOPHAN AMINOTRANSFERASE RELATED2 (TAR2)*, and the resulting effect on auxin levels is required to mediate the T6P-induced activation of storage processes. Our results suggest that auxin acts downstream of T6P to facilitate seed filling, thereby providing a salient example of how a metabolic signal governs the hormonal control of an integral phase transition in a crop plant.

## Introduction

The transition from early patterning into seed filling is an important phase change in developing seeds, ensuring seed survival and the nourishment of seedling growth upon germination. For this reason, plants have evolved a regulatory network to control seed filling, and carbohydrates appear to play a pivotal role in this process (Weber et al., 2005; Hills, 2004). Sucrose is thought to have a dual function in developing seeds as a nutrient sugar and as a signal molecule triggering storage-associated gene expression (Weber et al., 1998). Two decades ago, the invertase control hypothesis of seed development was formulated, suggesting that seed coat-borne invertases prevent the onset of storage processes in the early embryo by cleaving the incoming sucrose into hexoses (Weber et al., 1995a). When invertase activity declines, sucrose levels begin to rise and seed filling is initiated. However, relatively little is known about the perception and signaling of this metabolic switch.

T6P, the intermediate of trehalose biosynthesis, has been shown to be an essential signal metabolite in plants, linking growth and development to carbon metabolism (Lunn et al., 2006; Figueroa et al., 2016b). The sucrose–T6P nexus model postulates that T6P acts as a signal of sucrose availability, helping to maintain sucrose levels within a range that is appropriate for the developmental stage of the plant (Yadav et al., 2014). The particular importance of T6P for developmental transitions in plants is underlined by a growing number of growth processes known to be affected by directed modulation of T6P levels or by mutations of several T6P biosynthesis genes (Satoh-Nagasawa et al., 2006; Debast et al., 2011; Wahl et al., 2013). A striking example with respect to seed development involves the mutation of TREHALOSE 6-PHOSPHATE SYNTHASE 1 (TPS1) that catalyzes the formation of T6P from glucose 6-phosphate and uridine diphosphate glucose (UDPG) in Arabidopsis (Blazquez et al., 1998). Loss of TPS1 causes embryo abortion at the point where the embryo transitions from torpedo to early cotyledon stage (Eastmond et al., 2002), while the accumulation of storage proteins and lipids is compromised (Gómez et al., 2006). This, together with the resulting rise in sucrose concentration, supports the hypothesis that T6P signals seed filling processes by adjusting the consumption of maternally-delivered sucrose.

The small size of Arabidopsis seeds, however, presents practical difficulties in investigating how T6P participates in the regulation of seed filling. Here, we make use of the large size of pea seeds, allowing the easy preparation and compositional analysis of individual embryos. We engineered transgenic pea plants for embryo-specific expression of T6P synthase (TPS) and T6P phosphatase (TPP) genes from *Escherichia coli*, affecting T6P content and seed filling in parallel. Our results provide genetic and biochemical evidence that T6P reports the raising sucrose status in the maturing embryo, leading to a stimulation of embryo growth and reserve starch biosynthesis. Moreover, our findings show that auxin acts as a key mediator of this process.

## Results

### Cotyledon Differentiation and Starch Accumulation are Impaired by Embryo-specific Expression of TPP

To assess the potential role of T6P in the control of seed filling, we performed an initial metabolite analysis of growing pea embryos, revealing that T6P levels increased at the transition phase in parallel with sucrose and remained at high levels during the storage phase (Figure 1A). There was a positive correlation between T6P and sucrose (Pearson correlation coefficient, *r*=0.84), suggesting that T6P might control the phase transition into seed filling in response to sucrose accumulation. Next, we made use of a well-established approach to manipulate T6P levels in plants (Schluepmann et al., 2003), and reduced the T6P content in developing pea embryos by heterologous expression of a bacterial TPP, encoded by the *otsB* gene from *Escherichia coli*. These transgenic *USP::TPP* lines were derived from a set of 18 independent T_1_ plants, five of which were used to establish transgene homozygotes. The expression of the *USP::TPP* transgene was confirmed by quantifying the activity of TPP and the content of T6P in the developing embryos of these lines. While no TPP activity was detectable in WT embryos, considerable activity was present in *USP::TPP* embryos (Supplemental Table 1). This led to a significant depletion of the T6P content in the transgenic embryos (Supplemental Table 1), resulting in much smaller seeds and a wrinkled seed phenotype at maturity (Figure 1B, Supplemental Table 2). Mendel’s wrinkled-seed trait is associated with impaired reserve starch synthesis (Bhattacharyya et al., 1993), as was also the case for the seeds set by *USP::TPP* plants (Figure 1C), which contained 50% less starch on a per seed basis (Supplemental Table 2). At the same time, sucrose levels were elevated in transgenic embryos (Figure 1D), indicating that the sucrose-to-starch conversion was affected. Microscopic examination supported this finding, with *USP::TPP* embryos harboring considerably fewer and smaller starch granules than wild-type (WT) embryos (Figure 1E). However, altered starch accumulation only partially explains the diminished dry weight of *USP::TPP* seeds (Supplemental Table 2). Reduced cotyledon growth and smaller cell size also contribute to the decrease in dry weight (Figure 1F, Supplemental Table 3). These effects acted together to compromise the increase in embryo fresh weight at late stages of development (Figure 1G), while early embryo growth and organ determinacy were unaffected by the presence of the transgene (Supplemental Figure 1). Nuclear magnetic resonance (NMR) imaging of *USP::TPP* cotyledons revealed a substantial impairment in the formation of a spatial gradient in T_2_ transverse relaxation time (Figure 1H). The differences between the T_2_ signal of WT and transgenic embryos are mainly due to reduced enlargement of starch granules as well as increasing vacuolization towards the abaxial (inner) parts of the differentiating *USP::TPP* cotyledons (Van As, 2006; Borisjuk et al., 2012). Altogether, altered differentiation of *USP::TPP* embryos is in good agreement with defective cotyledon growth in the Arabidopsis *tps1* mutant (Eastmond et al., 2002; Gómez et al., 2006). These results indicate that T6P is a key factor in mediating the phase transition from patterning into seed filling, with at least two major processes being influenced by T6P: the conversion of sucrose into reserve starch, and the gradual differentiation of cotyledons.

**Figure 1.**
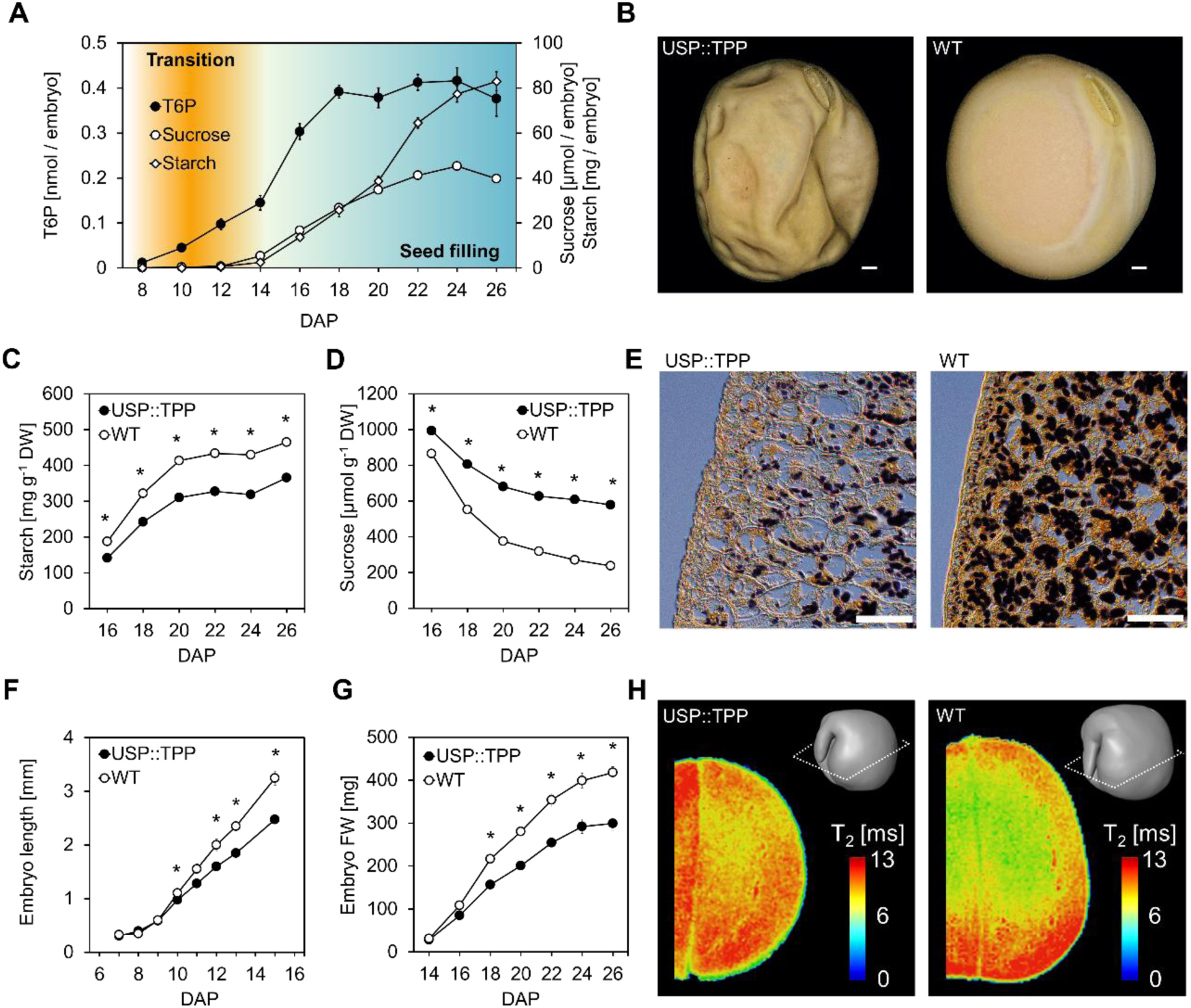
T6P promotes reserve starch accumulation and cotyledon differentiation in pea. (**A**) The relationship between the level of T6P and the accumulation of sucrose and starch in wild-type (WT) embryos over a period from 8 to 28 days after pollination (DAP). The error bars indicate the SEM (*n*=5). (**B**) The appearance of mature seeds formed by *USP::TPP* #3 and the corresponding WT plants. Scale bar, 1 cm. (**C**, **D**) Starch (C) and sucrose (D) levels in growing *USP::TPP* and WT embryos. Error bars indicate SEM (*n*=25), * *P* ≤ 0.001 (Student’s *t*-test). (***E*)** Iodine staining of starch granules in 22-day-old embryos harvested from *USP::TPP* #3 plants and corresponding WT plants. Scale bars, 500 µm. (**F**, **G**) Length (F) and fresh weight (G) of developing *USP::TPP* and WT embryos. Values given as means ± SEM (f: *n*=5, g: *n*=25), * *P* ≤ 0.05 (Student’s *t*-test). (**H**) Quantitative NMR imaging of transverse relaxation time (T_2_) in living *USP::TPP* #3 and WT cotyledons at 26 DAP. The 3D-scheme on the right indicates the virtual cross-section plane used for visualization. T_2_ values are color-coded.

### T6P Promotes Sucrose-to-Starch Conversion by Activating Key Enzymes of Starch Synthesis at the Transcript Level

Our current understanding of starch biosynthesis in pea seeds derives from extensive studies on mutations at different *rugosus* (*r*) loci, which curtail the activity of individual starch enzymes (Supplemental Figure 2A). In an attempt to identify the enzymatic steps within the sucrose-to-starch conversion process which were regulated by T6P, we compared the metabolic changes in *USP::TPP* embryos with those elicited by *r*, *rb*, and *rug4* mutations (Bhattacharyya et al., 1990; Hylton and Smith, 1992; Craig et al., 1999). In transgenic embryos, concentrations of hexose phosphates and UDPG were consistently elevated (Supplemental Figure 2B), while only adenosine diphosphate glucose (ADPG) levels were markedly lower than in WT (Figure 2A). A similar result was obtained only in the developing *rb* embryos (Supplemental Table 4), which have impaired ADPG-pyrophosphorylase (AGP) activity (Hylton and Smith, 1992), suggesting that the reduced starch accumulation in *USP::TPP* embryos is due to a similar defect. The peak level of AGP activity in transgenic embryos never rose to the height seen in the WT embryo, remaining some 68% lower during the main storage phase (Figure 2B). This, together with a constant decrease in total phosphoglucomutase (PGM) activity (Supplemental Figure 3A), largely explains the lower starch levels during the entire storage phase. To address the question of how T6P modulates the activity of these enzymes, we initially analyzed the degree of monomerization of the AGP small subunits as an estimate of the redox activation state, but no substantial difference was detected between transgenic and WT embryos (Supplemental Table 5). This is consistent with the recent finding that T6P has little or no effect on the post-translational modulation of AGP in Arabidopsis leaves (Martins et al., 2013). To establish whether any transcriptional regulation was involved, we monitored the transcript abundance of a number of genes encoding PGM and subunits of AGP, revealing that in the transgenic embryos, *PGM2*, *AGPL* (encoding the AGP large subunit), and the two small subunit encoding genes *AGPS1* and *AGPS2* were repressed (Figure 2C, Supplemental Figure 3B). The strongest reduction was recorded for *AGPL*, the transcription of which tends to peak during the period when the pea seed is most rapidly accumulating starch (Burgess et al., 1997). The loss of AGPL is the underlying cause for seed wrinkling in *rb* mutants (Hylton and Smith, 1992), suggesting that the wrinkled phenotype observed in *USP::TPP* seeds is due to a reduction in *AGPL* expression. The implication of these results is that T6P controls the conversion of sucrose into starch, at least in part, by modulating AGP activity at the transcript level.

**Figure 2.**
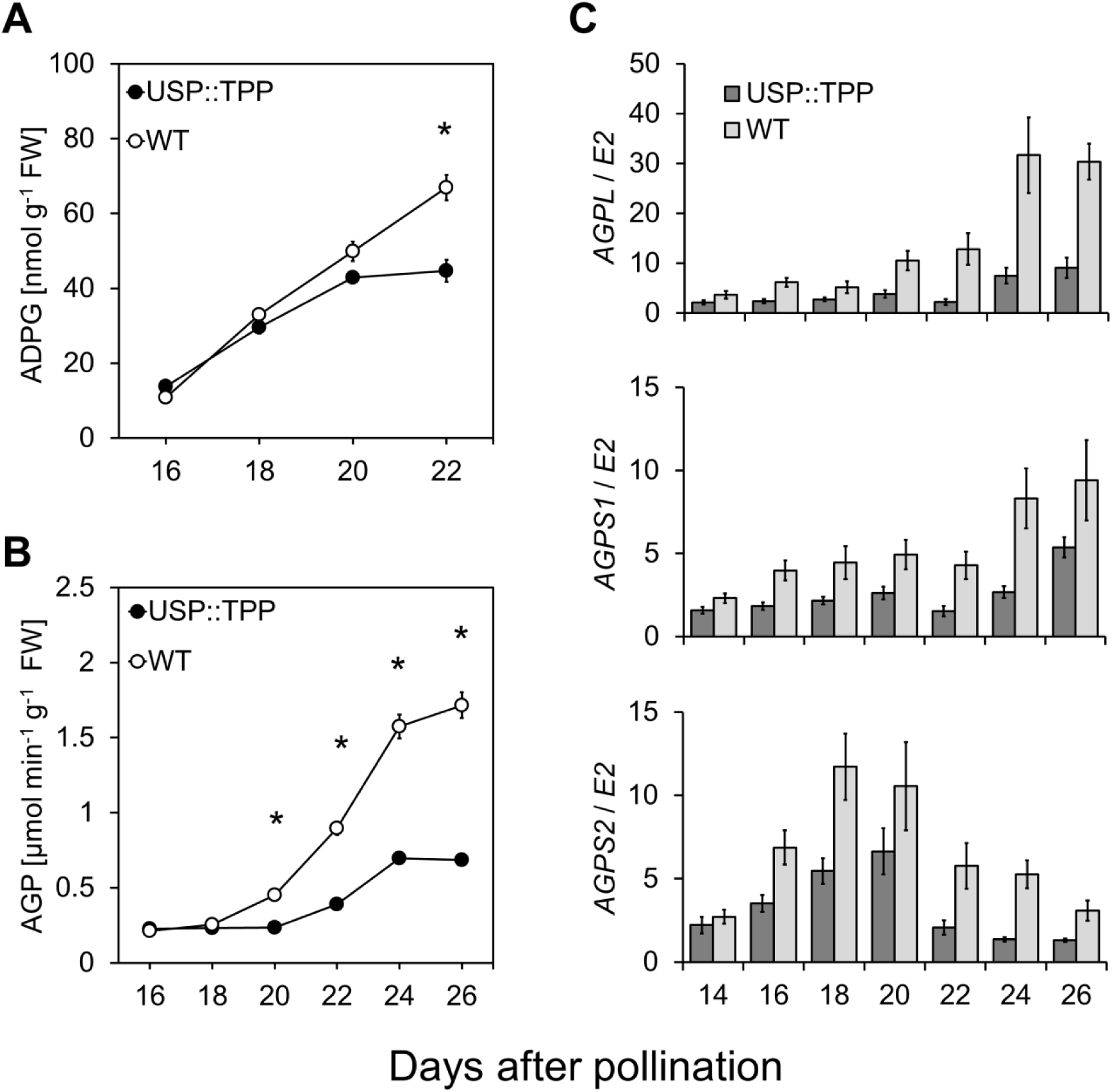
The effect of heterologous TPP expression on AGP activity and transcript levels of the corresponding genes. (**A**, **B**) The levels of ADPG (A) and AGP activity (B) in maturing *USP::TPP* and WT embryos. Values are means ± SEM (*n*=25), * *P* ≤ 0.001 (Student’s *t*-test). (**C**) Relative abundance of *AGPL*, *AGPS1* and *AGPS2* transcripts in *USP::TPP* and WT embryos over a period from 14 to 26 DAP. Values given as means ± SEM (*n*=10).

### Auxin Acts Downstream of T6P to Facilitate Seed Filling

To uncover novel signaling components that mediate the effects of T6P on seed filling, we performed a microarray analysis of embryos harvested at either the transition or main storage phase, focusing on genes that are consistently repressed in *USP::TPP* embryos. This analysis identified *TAR2*, the expression of which was unique in being dramatically reduced (Figure 3A), as a possible target of T6P. In pea, TAR2 participates in auxin biosynthesis via the indole-3-pyruvic acid pathway, and mutation of the corresponding gene leads to reduced levels of 4-chloro-indole-3-acetic acid (4-Cl-IAA), the predominant auxin in maturing seeds (Tivendale et al., 2012). Remarkably, the *tar2-1* mutation affects the phase transition into seed filling, resulting in the formation of small, wrinkled seeds with decreased starch content and a considerably lower level of AGP activity (McAdam et al., 2017). Our finding that *USP::TPP* seeds phenocopied those of the *tar2-1* mutant, and that introduction of *USP::TPP* into a *tar2-1* background had no additional phenotypic effects beyond those of the parental lines (Table 1, experiment 1 and Figure 3B), raises the possibility that both T6P and TAR2 act in the same signaling pathway. Measurement of the auxin content of *USP::TPP* embryos revealed a notable decrease in that of 4-Cl-IAA, by up to 70% (Figure 3C), while at the same time the content of the TAR2-specific substrate, 4-Cl-tryptophan, was higher than in WT embryos (Figure 3D). Together with the considerable increase of T6P in *tar2-1* embryos (Figure 3E), these results provide evidence that T6P acts as an upstream regulator of TAR2. As proof of this, we created hybrids between *USP::TPP* plants and transgenic plant lines harboring the *USP::TAR2* transgene, which directs expression of the *TAR2* coding sequence under the control of the embryo-specific *USP* promoter. To generate combinations of both transgenes, two previously generated *USP::TAR2* lines #3 and #5 (McAdam et al., 2017) were crossed with *USP::TPP* #2 and #3 plants (Table 1, experiment 2). Segregants from these crosses were used to establish three double transgene homozygotes, referred to as *USP::TPP* #2/*USP::TAR2* #3, *USP::TPP* #3/*USP::TAR2* #3 and *USP::TPP* #3/*USP::TAR2* #5. The activity of the *USP::TAR2* transgene was able to largely restore seed size and starch content to WT levels (Table 1, experiment 2, and Figure 3F), even though the level of TPP activity was still considerable (Supplemental Table 6). Regardless of these reconstituting effects, embryos formed by the homozygous *USP::TPP*/*USP::TAR2* plants shared the same pale green color as those formed by *USP::TPP* plants (Figure 3G), indicating that T6P regulates developmental processes in addition to those involving the TAR2 pathway. Taken together, our findings strongly suggest that normal cotyledon growth and reserve starch accumulation are both dependent on the transcriptional activation of *TAR2* by T6P.

**Figure 3.**
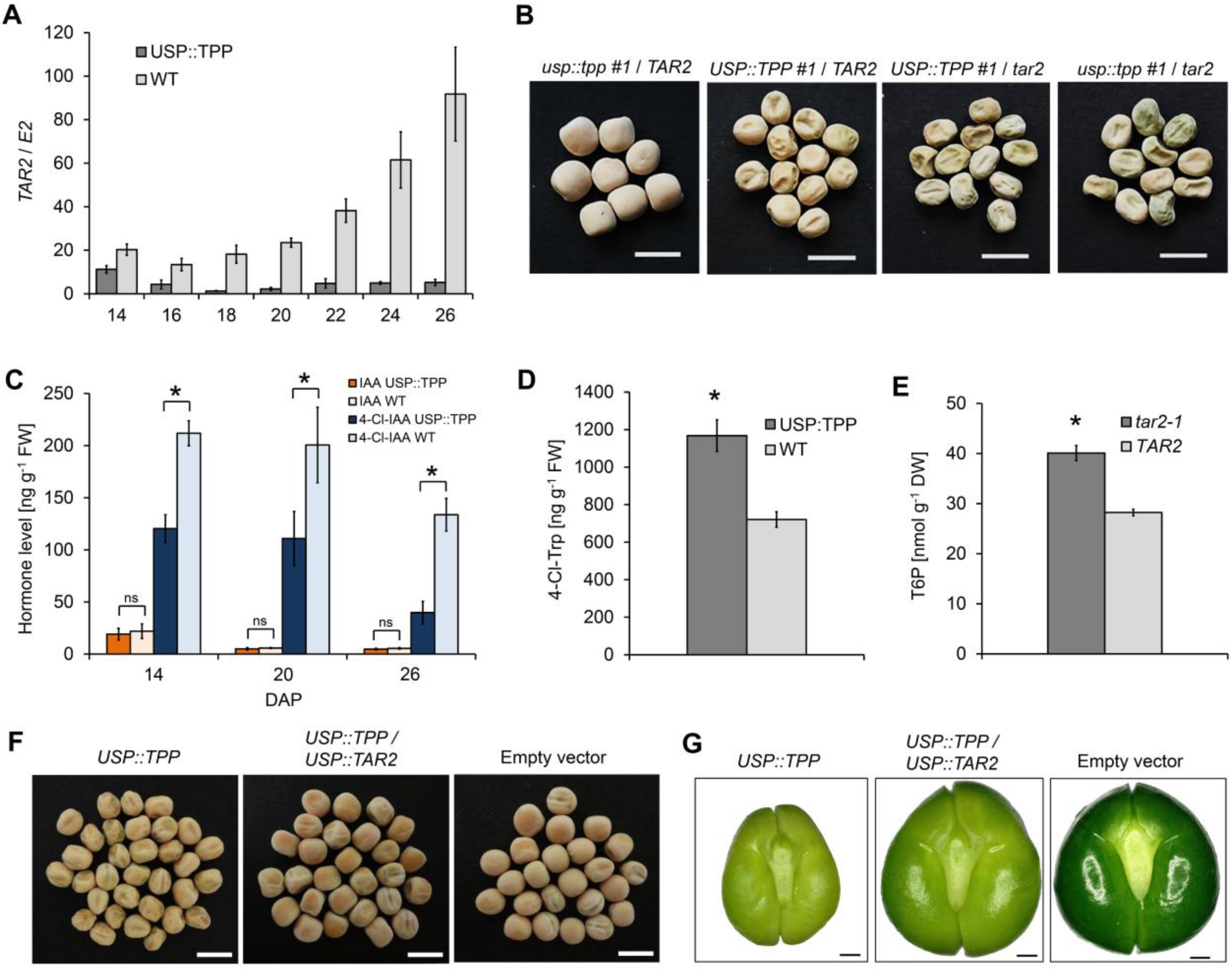
Expression of TPP affects auxin synthesis in developing pea embryos. (**A**) Relative abundance of *TAR2* transcripts in 14 to 26-day-old *USP::TPP* and WT embryos. Error bars denote upper and lower limit of SEM (*n*=10). (**B**) The photographs show dry seeds harvested from *usp::tpp* #1 / *TAR2*, *USP::TPP* #1 / *TAR2*, *USP::TPP* #1 */ tar2-1*, and *usp::tpp* #1 / *tar2-1* plants. Scale bar, 1 cm. (**C**) Auxin levels in growing *USP::TPP* and WT embryos. Values are means ± SEM (*n*=9), * *P* ≤ 0.05 (Student’s *t*-test), ns, not significant. (**D**) Levels of 4-Cl-tryptophan (4-Cl-Trp) in 26-day-old *USP::TPP* and WT embryos. Values are means ± SEM (n=9). Significant difference according to Student’s t-test: *P ≤ 0.05. (**E**) The content of T6P in 22-day old *tar2-1* and WT embryos. Values are means ± SEM (n=6), * *P* ≤ 0.001 (Student’s *t*-test) (**F**) The seed phenotype of hybrids between *USP::TPP* and *USP::TAR2* plants. The photographs present dry seeds from plants harboring *USP::TPP*#3, *USP::TPP*#3/*USP::TAR2*#3, and the empty vector control. Scale bars, 1 cm. (**G**) The appearance of 22-day-old embryos developing on *USP::TPP* #3, *USP::TPP* #3/*USP::TAR2* #3, and empty vector plants. Scale bars, 1 mm.

**Table 1.** The effect on seed weight and starch content by introducing either the *tar2-1* mutant allele (experiment 1) or the *USP::TAR2* transgene (experiment 2) into *USP::TPP* plants.

|  | Starch | Seed weight |
| --- | --- | --- |
| Experiment 1 | *mg g<sup>-1</sup> DW* | *mg* |
| *TAR2* | 438 ± 8 a | 335 ± 23 a |
| *tar2-1* | 311 ± 16 b | 155 ± 10 c |
| Empty vector / *TAR2* | 422 ± 15 a | 323 ± 11 a |
| Empty vector / *tar2-1* | 340 ± 17 b | 129 ± 9 c |
| *USP::TPP* #1 / *TAR2* | 356 ± 10 b | 212 ± 7 b |
| *USP::TPP* #1 / *tar2-1* | 348 ± 3 b | 148 ± 7 c |
| *usp::tpp* #1 / *TAR2* | 433 ± 12 a | 313 ± 10 a |
| *usp::tpp* #1 / *tar2-1* | 336 ± 13 b | 139 ± 7 c |
| *USP::TPP* #2 / *TAR2* | 350 ± 10 b | 203 ± 11 b |
| *USP::TPP* #2 / *tar2-1* | 349 ± 9 b | 151 ± 6 c |
| *usp::tpp* #2 / *TAR2* | 429 ± 10 a | 312 ± 8 a |
| *usp::tpp* #2 / *tar2-1* | 342 ± 5 b | 150 ± 7 c |
| Experiment 2 |  |  |
| Empty vector | 497 ± 12 a | 298 ± 3 a |
| *USP::TPP* #2 | 397 ± 23 b | 193 ± 14 b |
| *USP::TPP* #3 | 384 ± 8 b | 199 ± 12 b |
| *USP::TAR2* #3 | 466 ± 22 a | 320 ± 8 a |
| *USP::TAR2* #5 | 521 ± 23 a | 306 ± 6 a |
| *USP::TPP* #2 / *USP::TAR2* #3 | 507 ± 13 a | 306 ± 18 a |
| *USP::TPP* #3 / *USP::TAR2* #3 | 482 ± 21 a | 277 ± 5 a |
| *USP::TPP* #3 / *USP::TAR2* #5 | 497 ± 30 a | 312 ± 7 a |

Values are means ± SEM (n=5). Means labeled with the same letter (a-c) do not differ from one another ( $P \leq 0.01$ ).

### Embryo-Specific Elevation of T6P Induces Auxin and Starch Biosynthesis

We next investigated the effect of elevated T6P on seed filling processes by heterologously expressing *otsA*, an *Escherichia coli* gene encoding TPS, in an embryo-specific manner. To this end, five homozygous *USP::TPS* lines were generated from a set of 22 independent T_1_ plants. Analysis of developing embryos harvested from three of these lines showed that they contained considerably more T6P than those of their sibling WT embryos (Supplemental Table 7), confirming the functional expression of the bacterial TPS. Consistent with similar experiments conducted in both Arabidopsis and potato (Yadav et al., 2014; Debast et al., 2011; Schluepmann et al., 2003), the activity of the transgene resulted in a substantial depletion in the content of soluble sugars (Supplemental Figure 4). Short-term, induced elevation of T6P in Arabidopsis has been shown to induce a loss in sucrose content, due to a shift in assimilate partitioning away from sucrose in favor of organic and amino acids in the light, or inhibition of transitory starch turnover in the dark (Martins et al., 2013; Figueroa et al., 2016a). The exposure of wheat plants to cell-permeable forms of T6P has been shown to promote the size and starch content of the grain, and to raise AGP activity (Griffiths et al., 2016), an observation which ties in with the increased AGP activity (Figure 4A) in TPS expressing pea embryos. Compared to WT, expression of *TAR2* was induced in *USP::TPS* expressing embryos (Figure 4B) accompanied by an increase in 4-Cl-IAA levels at later stages (Figure 4C). It appears that the elevation of T6P has a positive influence on the sucrose-to-starch conversion by inducing AGP, and we conclude that this is mediated by a prolonged stimulation of auxin synthesis via TAR2. Despite these favorable changes, neither the starch content nor the size of *USP::TPS* seeds was affected (Supplemental Table 8). This may not be surprising, considering that the limits to the final size of the embryo and its capacity to accumulate dry matter are largely influenced by the maternal genotype and assimilate supply from the seed coat (Davies, 1975; Weber et al., 1996).

**Figure 4.**
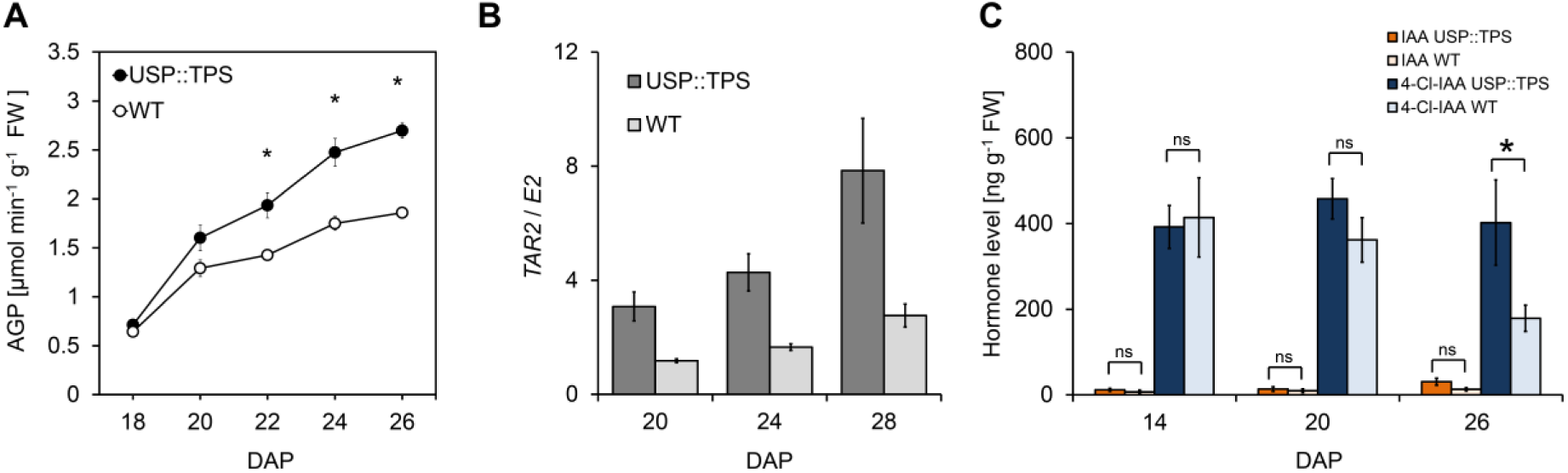
Expression of TPS induces starch and auxin synthesis in developing pea embryos. (**A**) The level of AGP activity in 18- to 26-day-old *USP::TPS* and WT embryos (*n=*15), * *P* ≤ 0.05 (Student’s *t*-test). (**B**) Relative transcript levels of *TAR2* in embryos formed by *USP::TPS* and corresponding WT plants. Transcript abundances are means ± SEM (*n*=8). (**C**) Auxin accumulation in growing *USP::TPS* and WT embryos. Values are means ± SEM (*n=*9),* *P* ≤ 0.05 (Student’s *t*-test), ns; not significant.

## Discussion

Efficient deposition of storage compounds in seeds is a key determinant of crop yield. The interplay between carbohydrates and hormones seems to play a crucial role in the control of seed filling, but the underlying regulatory network of this process remains undefined. In this study, we provide several lines of evidence showing interaction between the signaling sugar T6P and the major plant hormone auxin, as a requisite for normal seed filling in pea. We showed that T6P and sucrose levels increased in parallel at the time point when the embryo starts to build up storage products, and by manipulating the T6P content in embryos, we found that the transition into the storage mode is based on this relationship. Our results imply that T6P regulates seed filling by promoting cell differentiation and starch accumulation in the maturing embryo, thereby allowing efficient utilization of incoming sucrose. This finding is in agreement with the T6P-sucrose nexus (Yadav et al., 2014) and complements the existing view on the dual function of sucrose as a key metabolite and signaling molecule of seed filling (Weber et al., 2005; Hills, 2004), with T6P reporting the change in sucrose concentration to the regulatory network of embryo differentiation. Like in most sink organs, maturing embryos receive sucrose from the phloem and its cleavage is the initial step in the direction of storage product synthesis. However, sucrose is also required to induce storage-related gene expression causing upregulation of important enzymes like AGP (Müller-Röber et al., 1990; Weber et al., 1998). Furthermore, cell expansion in explanted *Vicia faba* embryos is triggered in response to sucrose feeding (Weber et al., 1996). The evidence presented here clearly indicate that most effects which previously have been ascribed to a signaling function of sucrose are principally controlled via a T6P-mediated pathway. We suggest that T6P connects the sucrose state with other regulatory components involved in the control of storage metabolism and embryo differentiation, such as SnF1-related protein kinase1 (Radchuk et al., 2006). This energy sensor coordinates metabolic and hormonal signals with embryo growth (Radchuk et al., 2010). In developing tissues, SnRK1 activity is inhibited by T6P in the presence of a so far uncharacterized protein (Zhang et al., 2009), and binding of T6P to the catalytic subunit (SnRK1α1) disrupts association and activation of SnRK1 by the SnRK1 activating kinase (SnAK)/Rep-Interacting Kinase1 (GRIK) protein kinases (Zhai et al., 2018).

Until now, the underlying mechanism by which T6P integrates carbohydrate partitioning with the hormonal control of plant development has not been apparent. Importantly, our data now indicate that auxin is a key factor in mediating the effects of T6P, which acts upstream of the pivotal auxin biosynthesis gene *TAR2* (McAdam et al., 2017) to trigger seed filling in pea. We propose that this process is mediated via a modulation of the auxin 4-Cl-IAA (Figure 5), the concentration of which increases sharply at the transition stage (Tivendale et al., 2012). There is a growing body of evidence that soluble sugars control plant growth by modifying auxin biosynthesis (LeClere et al., 2010; Sairanen et al., 2012; Lilley et al., 2012; Barbier et al., 2015). Altered sugar concentrations in endosperm-defective *miniature1* kernels of maize have been suggested to induce auxin deficiency due to the suppression of the genes *ZmTAR1* and *ZmYUCCA1* (LeClere et al., 2010). Both of these genes encode proteins involved in the indole 3-pyruvic acid branch of auxin biosynthesis (Won et al., 2011; Stepanova et al., 2011), with *ZmYUCCA1* being essential for the formation of a normal endosperm (Bernardi et al., 2012). Apart from seeds, a similar connection between sugars and auxin has been implicated in the control of shoot branching. Contradicting the classical theory of apical dominance (Thimann et al., 1934), the accumulation of sucrose enables the initiation of bud outgrowth after decapitation in pea (Mason et al. 2014), an effect thought to be mediated by T6P (Fichtner et al., 2017). Notably, feeding of sucrose stimulates auxin synthesis within buds and promotes sustained auxin export from bud to stem (Barbier et al, 2015). Collectively, our findings indicate that the hitherto unknown interaction between T6P and auxin might play a general role in mediating the sugar-auxin link. Of ongoing interest will be to determine how this relationship fits within the current understanding of the regulatory frameworks surrounding growth processes and developmental transitions in plants.

**Figure 5.**
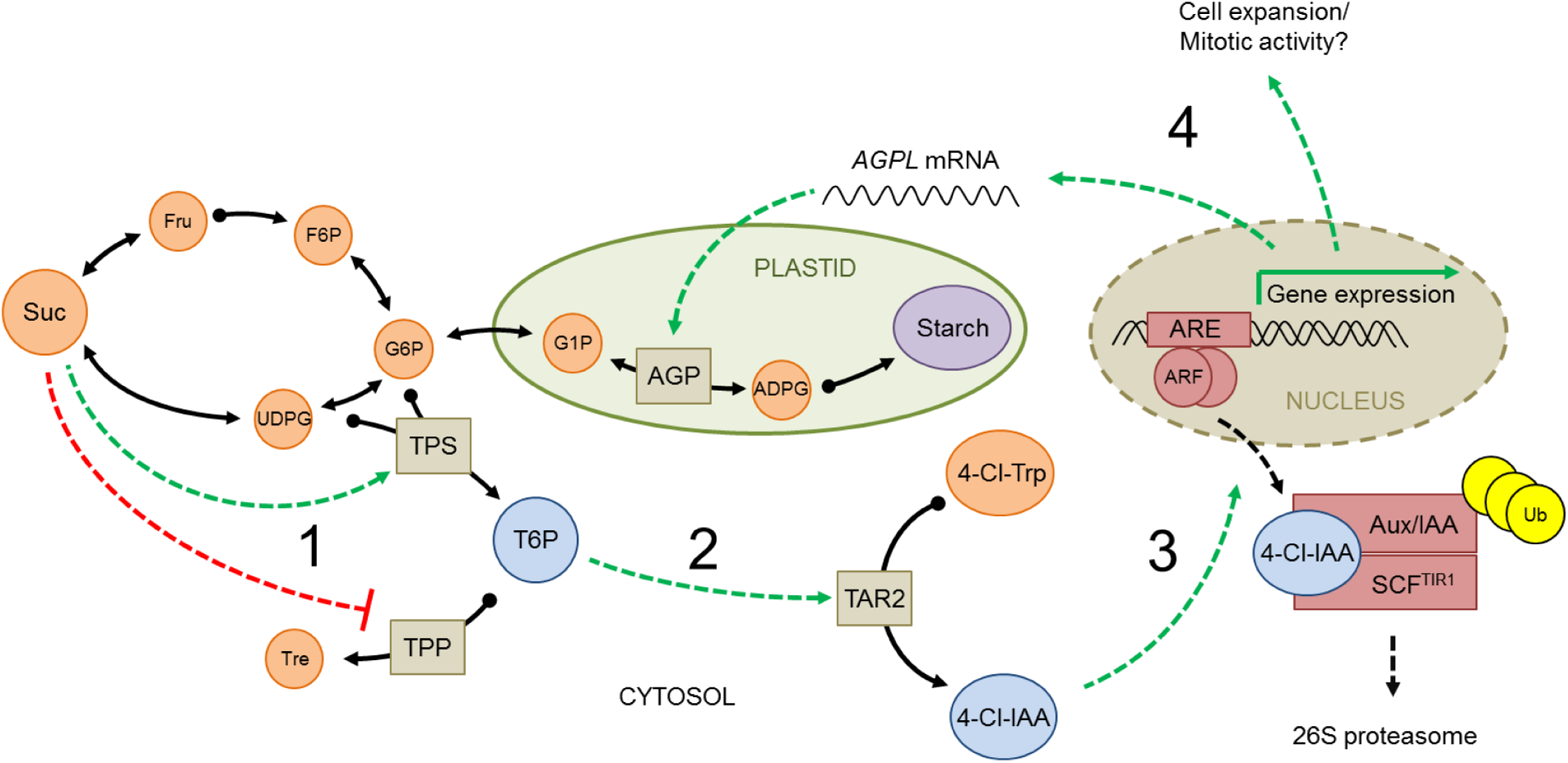
A simplified model of the T6P-auxin signaling pathway regulating embryo maturation in pea. (*1*) During the transition from early pattern formation to seed filling, maternally delivered sucrose accumulates in the embryo, raising the level of T6P. (*2*) Activation of T6P signaling is required for the expression of *TAR2* and for the increased synthesis of 4-Cl-IAA from 4-Cl-Trp. (*3*) A rise in the level of 4-Cl-IAA derepresses auxin responsive genes by promoting the ubiquitin-mediated release of the AUX/IAA repressor from ARF via the activation of the Aux/IAA-SCF^TIR1^ co-receptor system. (*4*) The transcriptional activation of starch synthesis genes, in particular *AGPL*, is necessary for normal starch accumulation, while certain as yet unidentified target genes regulate cotyledon growth via the stimulation of cell proliferation. Together, these processes act to efficiently allocate the incoming sucrose within the differentiating embryo, and to ensure continuous growth and optimal filling of the maturing seed. 4-Cl-IAA, 4-Cl-indole-3-acetic acid; 4-Cl-Trp, 4-Cl-tryptophan; ADPG, ADP-glucose; AGP, ADPG pyrophosphorylase; ARE, auxin responsive element; ARF, auxin response factor; F6P, fructose 6-phosphate; Fru, fructose; G1P, glucose 1-phosphate; G6P, glucose 6-phosphate; Suc, sucrose; T6P, trehalose 6-phosphate; TPP, T6P phosphatase, TPS, T6P synthase; Tre, trehalose; Ub, ubiquitin.

## Methods

### Plant material

Transgenic pea plants were created within the cv. ‘Erbi’, previously described for transgenic *USP::TAR2* plants (McAdam et al., 2017). The *tar2-1* mutant was made in the background of cv. ‘Cameor’ (Tivendale et al., 2012), while those carrying non WT alleles at *r*, *rb*, and *rug4* were near isogenic selections made in, respectively, germplasm accessions WL 200, WL 1685 and SIM91 (John Innes Germplasm Collection, Norwich, UK) (Wang and Hedley, 1993). Plants were grown under a 16 h photoperiod provided by artificial light (550 µmol m^−2^ s^−1^). The light/dark temperature regime was 19°C/16°C. The plants were fertilized once a week with 0.4% Hakaphos® blau 15+10+15(+2) (Compo Expert, Münster, Germany) starting four weeks after sowing. Flowers were tagged at the time of pollination, and seeds were harvested around midday according to the number of days after pollination (DAP) which had elapsed. Embryos were excised from 2-3 seeds per pod, weighed and snap-frozen in liquid nitrogen.

### Transgene construction and the production of transgenic plants

To generate transgenic *USP::TPS* and *USP::TPP* pea plants, coding sequences of the respective *Escherichia coli* genes *otsA* and *otsB* were PCR amplified from plasmids harboring the corresponding cDNAs. The oligonucleotides used to attach an *Xba*I restriction site to each end of the amplified coding sequences are listed in Table S9. The resulting amplicons were *Xba*I restricted and ligated into the *USP::pBar* binary vector which contains an embryo-specific expression cassette based on the long version of the *USP* promoter (Zakharov et al., 2004). Selected plasmids were sequenced for validation purposes and then introduced into *Agrobacterium tumefaciens* strain EHA 105. The generation of transgenic pea plants was performed according to a modified transformation method using sections from embryo axis (Schroeder et al., 1993). For this purpose, embryo axes were excised from germinating pea seeds (3 days after imbibition), sliced longitudinally into five to seven segments with a scalpel blade, and the obtained explants were immersed in a suspension of Agrobacteria. After two days of cocultivation on B_5_h medium (Brown and Atanassov, 1985), explants were washed with sterile water and transferred to selective P1 medium (Schroeder et al., 1993) containing 10 mg L^−1^ phosphinothricin (Duchefa, Haarlem, The Netherlands). After two weeks of callus formation, shoot growth was induced by cultivation on MS4 medium containing Murashige and Skoog macro and micronutrients (Murashige and Skoog, 1962), B5 vitamins (Gamborg et al., 1968), 4 mg L^−1^ 6-benzylaminopurine, 2 mg L^−1^ naphthalene acetic acid, 0.1 mg L^−1^ indol-3-yl butyric acid, 3% (w/v) sucrose supplemented with 10 mg L^−1^ phosphinothricin. When the developing shoots were about 5 cm in length, a substantial gain in plant growth was induced by grafting the shoots onto a WT root stock. The grafted plantlets were potted and maintained in growth chambers, where they subsequently flowered and produced seeds. Insertion and segregation of the transgenes was verified by PCR using oligonucleotides that are listed in Table S9. The allelic status at the *TAR2* locus was assessed by previously described PCR-based genotyping (Tivendale et al., 2012).

### Determination of sucrose, starch, total carbon and nitrogen

Snap-frozen embryos were ground to powder and lyophilized at −20°C. Mature seeds were pulverized in a ball mill and the powder dried in a desiccator. To measure tissue sucrose and starch contents, the powder was extracted twice in 80% (v/v) ethanol at 60°C and the supernatants pooled and vacuum-evaporated; the residue was dissolved in sterile water. Sucrose contents were determined enzymatically (Heim et al., 1993). The starch retained in the water-insoluble fraction was solubilized in 1 M KOH and gelatinized by incubating for 1 h at 95°C, after which it was neutralized by the addition of 1 M HCl. The starch content was determined as glucose units, following its complete hydrolysis to glucose using amyloglucosidase (Rolletschek et al., 2002). The carbon and nitrogen content of powdered seed tissue was obtained using a Vario Micro Cube elemental analyser (Elementar UK Ltd., Stockport, Great Britain).

### Enzyme activity assays

Enzyme activities were determined in growing embryos of three *USP::TPS*, five *USP::TPP*, and the corresponding WT lines each with five biological replicates per time point. The frozen, pulverized tissue was extracted in 5 vol. of 0.1 M MOPS (pH 7.4), 10 mM MgCl_2_, 1 mM EDTA, 1 mM EGTA and 2 mM DTT; the resulting homogenates were centrifuged (10,000×*g*, 4°C, 5 min) and the supernatants held on ice. The extracts were assayed for AGP (Weber et al., 1995b) and PGM (Manjunath et al., 1998) activity. TPP activity was determined by following the release of orthophosphate from trehalose 6-phosphate. The reaction mixtures, containing 25 mM HEPES/KOH, pH 7.0, 8 mM MgCl_2_, 0.05 mM Triton X-100, 0.5 mM EDTA, 1.25 mM T6P, and 2 µl of crude extract in a total volume of 20 µl, were incubated for 20 min at 32°C and stopped by applying 10 µl of 0.5 M HCl. The yield of orthophosphate was determined by adding 50 µL of 1% (w/v) ammonium molybdate dissolved in 1 M H_2_SO_4_ and 20 µL of 10% (w/v) ascorbic acid (Ames, 1966). After incubation at 40°C for 40 min, absorbance was immediately measured at 800 nm.

### AGP redox activation

The redox activation of the AGP of developing embryos was determined by monitoring the degree of monomerization of small AGP subunits in non-reducing SDS gels (Hendriks et al., 2003).

### RNA extraction, cDNA synthesis and transcript profiling

RNA was isolated from the frozen, pulverized tissue using a phenol/chloroform-based extraction method, followed by LiCl precipitation (Miranda et al., 2001). Contaminating genomic DNA was removed by incubating a 20 µg aliquot of the RNA in a 100 µL reaction containing 4 U TURBO DNA-free™ (Ambion/Life Technologies, Darmstadt, Germany), following the manufacturer’s protocol. After digestion, samples were desalted and concentrated to a volume of 20 µl by using Vivaspin 500 centrifugal concentrators with a molecular weight cut-off of 30,000 Da (Sartorius, Goettingen, Germany). The absence of genomic DNA contamination was confirmed by running a quantitative real time PCR (qRT-PCR) assay directed by a primer targeting intron #8 of the pea *PHOSPHOLIPASE C* (*PLC*) gene (Table S9). First strand cDNA was synthesized from 4.5 µg purified RNA using SuperScript III (Invitrogen, Carlsbad, USA) primed by oligo-dT, according to the manufacturer’s instructions. The reference sequence was a fragment of the pea ubiquitin-conjugating enzyme gene *E2* (PSC34G03; http://apex.ipk-gatersleben.de), whose expression stability in the developing pea seed was evaluated using geNorm software (Vandesompele et al., 2002). Profiling of *TAR2*, *AGPL*, *APGS1*, and *AGPS2* transcripts via qRT-PCR was performed as 10 µL reactions containing 2 µL of each primer (0.5 µM, sequences given in Table S9), 1 µL cDNA (1 µg/µl) and 5 µL Power SYBR**®** Green-PCR Master Mix (Applied Biosystems/Life Technologies, Darmstadt, Germany): the amplification regime consisted of a 95°C/10 min denaturation step, followed by 40 cycles of 95°C/15 s, 60°C/60 s. PCR amplification efficiencies were estimated using a linear regression method implemented in the LinRegPCR program (Ramakers et al., 2003). Relative transcript abundances were calculated using the 2^−ΔΔCt^ method (Livak and Schmittgen, 2001). For the qRT-PCR analysis of transgenic embryos, samples were obtained from two biological replicates per each of five *USP::TPP* and the corresponding WT lines, whereas four biological replicates each were sampled for two *USP::TPS* and the corresponding WT lines.

### Microarray hybridization and analysis

An 8×60K customized pea eArray (ID 045803, Agilent Technologies, Santa Clara, CA, USA) was used to scan the pea embryo transcriptome. Embryos were sampled at both 14 DAP and 22 DAP from two independent *USP::TPP* transgenic lines (three independent plants per line), along with 14 DAP embryos from six WT plants and 22 DAP embryos from five WT plants. Total RNA extracts were treated with RNase-free DNase and purified using the RNeasy RNA Isolation kit (Qiagen, Germany). A 100 ng aliquot of RNA was used to generate cRNA, which was Cy3 labelled via the Low Input Quick Amp Labeling Kit (Agilent Technologies, Santa Clara, CA, USA). The labelling efficiency, amount, as well as the amount and quality of the cRNA synthesized were monitored using an ND-1000 spectrophotometer (NanoDrop Technologies, Wilmington, DE, USA) and a Bioanalyser 2100 (Agilent Technologies, Santa Clara, CA, USA). A 600 ng aliquot of labelled cRNA was used for fragmentation and array loading (Gene Expression Hybridisation Kit, Agilent Technologies). Hybridization, scanning, image evaluation, and feature extraction were achieved as described by Pielot et al., 2015. The data were evaluated with the aid of Genespring v12.5 software (Agilent Technologies) using default parameters: relative expression values were achieved after log_2_ transformation, quantile normalization, and baseline transformation to the median of all samples. After removing outliers and transcripts without significant expression (absolute values ≥ 100), moderated *t*-test and false discovery rate (FDR) correction (Benjamini–Hochberg) were performed. Only features associated with a signal intensity difference of at least two fold between the WT and transgenic lines at a given sampling time point were retained. A high-stringency P cutoff (P *corrected≤*0.001) was used to remove random effects.

### Extraction and quantification of metabolites and phytohormones

Metabolites (including T6P) were extracted from the frozen, pulverized embryos of three *USP::TPS* and three *USP::TPP* transgenics, along with their corresponding WT sibling plants (five embryos per genotype at each time point). The content of soluble metabolites was assessed using a high-performance anion-exchange liquid chromatography coupled to tandem mass spectrometry (LC-MS/MS) (Figueroa et al., 2016a). To estimate the content of T6P in *tar2-1* and *TAR2* embryos, freeze-dried samples were extracted (Schwender et al., 2015) followed by ion chromatography using Dionex ICS-5000+ HPIC system (Thermo Scientific, Dreieich, Germany), coupled to a QExactive Plus hybrid quadrupol-orbitrap mass spectrometer (Thermo Scientific) equipped with a heated electrospray ionization probe. Chromatographic separation was performed on Dionex™ IonPac™ 2×50 mm and 2×250 mm AS11-HC-4 µm columns equilibrated with 10 mM KOH at 0.35 ml min^−1^ flow rate and 35°C column temperature. A linear gradient of 10–100 mM KOH was generated in 28 min followed by 2 min of column equilibration. The MS spectra were acquired using a full-scan range 67-1000 (m/z) in the negative mode at 140.000 resolving power, 200 ms maximum injection time and automated gain control at 1e6 ions. The source settings included 36 sheath gas flow rate, 5 auxiliary gas flow rate, 3.5 kV spray voltage. The capillary temperature was set to 320°C and S-lens was set to 50. Quantification was performed with external calibration using authenticated standard and TraceFinder 4.1 software package (Thermo Scientific). Auxins and 4-Cl-tryptophan were extracted from developing embryos harvested from three *USP::TPS* and three *USP::TPP* transgenic plants, and also from their corresponding sibling WT embryos, and subsequently quantified (three embryos per genotype at each time point) using ultra-performance liquid chromatography coupled with mass spectrometry (Tivendale et al., 2012).

### Histological and morphological analysis

Seeds were sliced into two pieces and fixed at room temperature overnight in 50% (v/v) ethanol, 5% (v/v) glacial acetic acid and 4% (v/v) formaldehyde. After dehydration by passing through an ethanol series, the samples were embedded in Paraplast Plus (Sigma-Aldrich, St. Louis, MO, USA). Cross sections of thickness 15 µm were mounted on poly-L-lysine-treated slides (Sigma-Aldrich) and stained with iodine in order to visualize the starch grains. The specimens were imaged by differential interference contrast (DIC) microscopy (Zeiss Axio Imager.M2), and Axiovision (Zeiss, Germany) software was used for scaling; cell areas were estimated from digital images using ImageJ software (/imagej.nih.gov/ij/), each measurement was based on sections from three biological replicates. Early embryo growth was verified by using a VHX digital microscope (Keyence, Osaka, Japan).

### Nuclear magnetic resonance imaging

Magnetic resonance experiments were performed by using a Bruker Avance III HD 400 MHz NMR spectrometer (Bruker BioSpin, Rheinstetten, Germany) equipped with a 1000 mT/m gradient system. For the measurement of the transverse relaxation time (T_2_) in embryonic tissues, excised pea embryos were placed in a saddle coil with an inner diameter of 10 mm. A standard Multi-Slice Multi-Echo sequence was applied in 3D (repetition time, 2500 ms; echo time, 5 ms; number of echoes, 12). The images were acquired with a resolution of 80 µm x 90 µm x 90 µm. The resulting datasets were processed by using the MATLAB software (MathWorks, Natick, MA, USA) with an in-house written algorithm. Calculation of the 3D T_2_-maps is based on a least-squares algorithm, and the resulting T_2_-maps were subsequently exported to AMIRA (FEI Visualization Sciences Group, Mérignac, France) for image processing.

### Data availability

Microarray data reported in this study have been deposited in the ArrayExpress database at EMBL-EBI (www.ebi.ac.uk/arrayexpress) under accession number E-MTAB-6659.

## Acknowledgments

We are grateful to Angela Schwarz, Elsa Fessel, Angela Stegmann and Katrin Blaschek for excellent technical assistance, and Noel Davies and David Nichols (Central Science Laboratory, University of Tasmania) for assistance with auxin analyses. We thank Mike Ambrose (John Innes Centre) for providing seeds, and Marion Dalmais, Richard Thompson, and co-workers (INRA, centre de Dijon) for isolating the *tar2-1* mutant. We also thank Bruno Müller, Hans Weber, and Winfriede Weschke for discussions and continuous support. This work was supported by Deutsche Forschungsgemeinschaft grants WE1641/16-1, GE878/5-2, and GE878/8-1.

## Author Contributions

T.M. directed the overall study design and performed enzyme activity measurements, microscopy, transcript profiling, compositional analysis of seeds, molecular cloning, plant transformation, cross breeding and genotyping. A.H., R.F., and J.E.L. conducted the LC-MS-based profiling of metabolites. I.T. and P.G. analysed the redox status of AGP. Microarray experiments were performed and analysed by R.R.. E.L.M. and J.J.R. performed the measurement of hormones. Nuclear magnetic resonance images were generated by E.M. and L.B.. T.M. wrote the paper, on which all authors commented.

## Supplementary Information

**Supplemental Figure 1.**
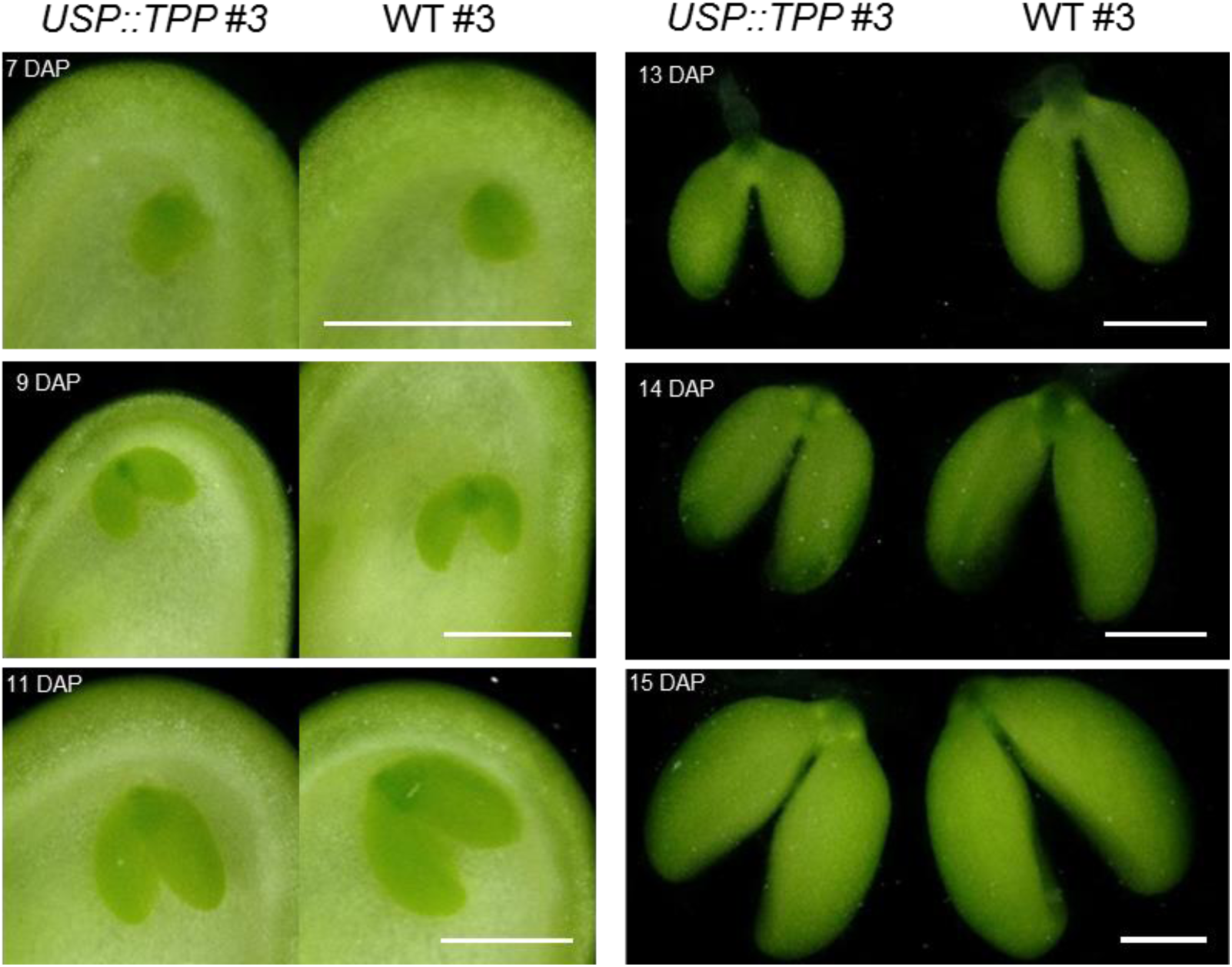
Early development of *USP::TPP* embryos. The photographs present embryos formed by the transgenic line *USP::TPP*#3 and WT#3, harvested at 7, 9, 11, 13, 14, and 15 days after pollination (DAP). Note the reduced length of transgenic cotyledons in embryos harvested later than 11 DAP. Scale bars: 1 mm.

**Supplemental Figure 2.**
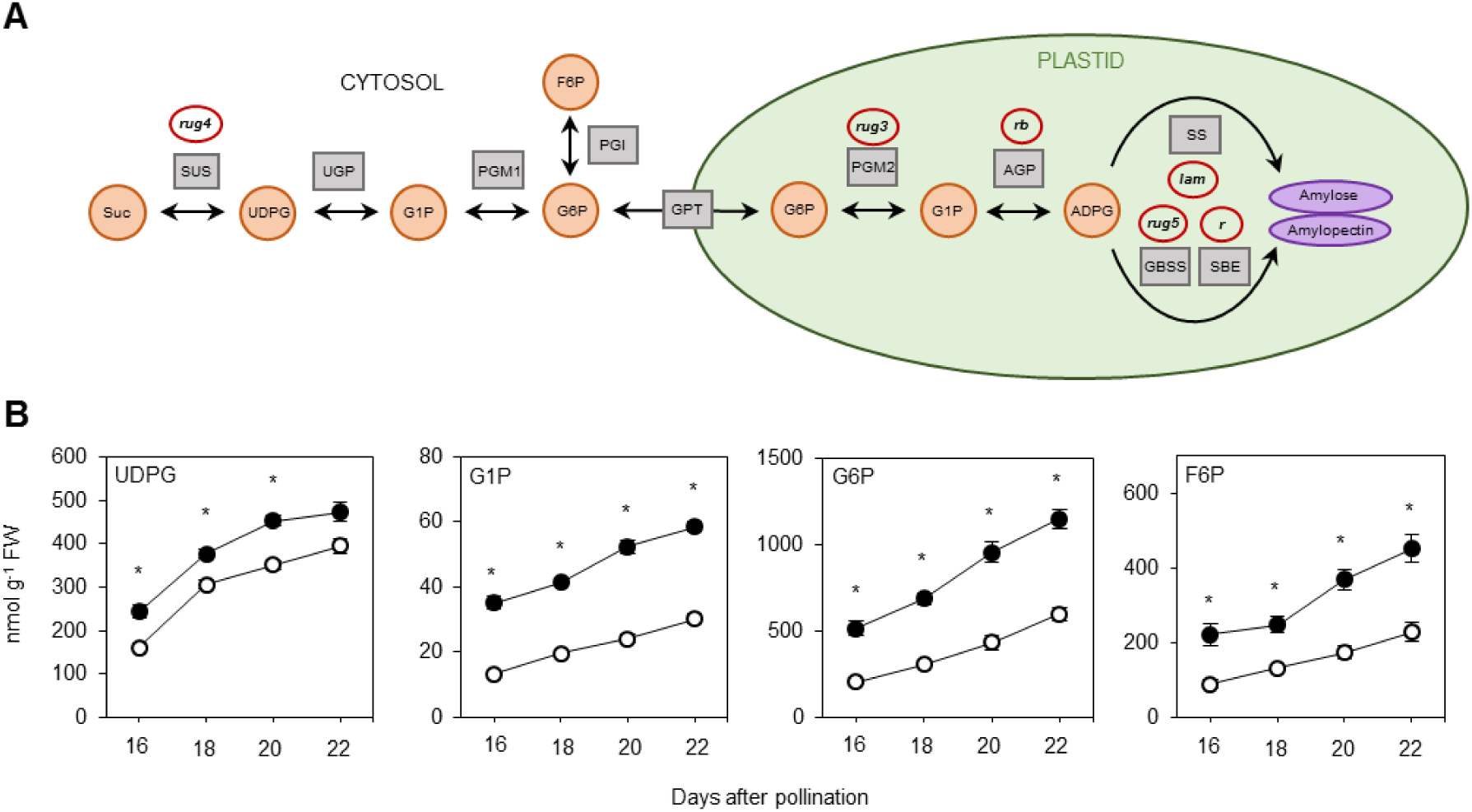
Analysis of phosphorylated intermediates directly involved in the sucrose-to-starch conversion. (**A**) Overview of the starch biosynthetic pathway in storage cells of pea embryos. Squares and circles symbolize enzyme activities and metabolites, respectively. Mutations affecting the corresponding enzyme activities are indicated by red rings. (**B**) Soluble sugar levels of developing *USP::TPP* (solid circles) and WT (open circles) embryos at 16, 18, 20, and 22 DAP. Error bars, SEM (n=25); significance was calculated according to Student’s t-test: *P ≤ 0.001. ADPG, ADP-glucose; AGP, ADP-glucose pyrophosphorylase; F6P, fructose 6-phosphate; G1P, glucose 1-phosphate; G6P, glucose 6-phosphate; GBSS, granule-bound starch synthase; GPT, glucose 6-phosphate/phosphate translocator; PGI, phosphoglucoisomerase; PGM, phosphoglucomutase; SBE, starch branching enzyme; Suc, sucrose; SUS, sucrose synthase; SS, starch synthase; UDPG, UDP-glucose; UGP, UDP-glucose pyrophosphorylase.

**Supplemental Figure 3.**
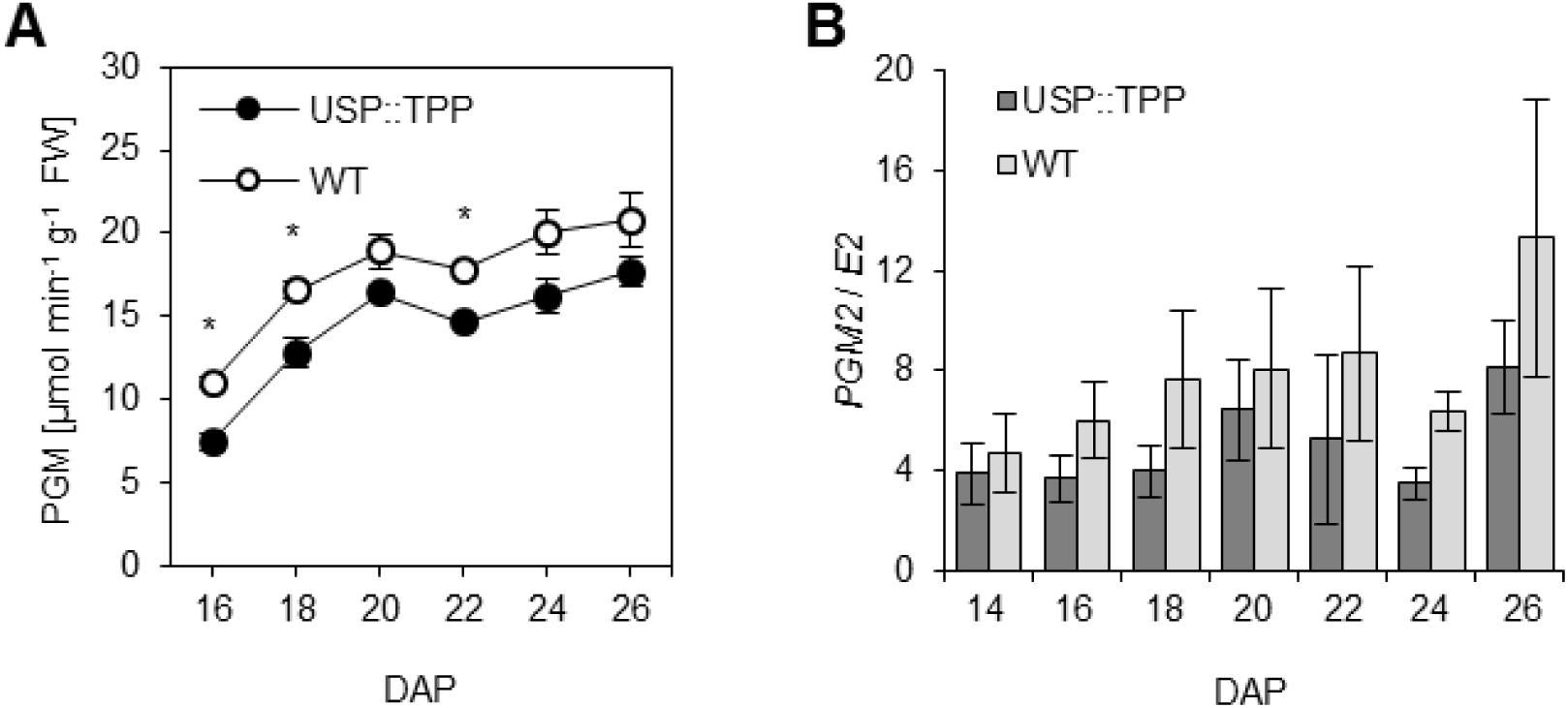
The effect of heterologous TPP expression on PGM activity. (**A**) Level of total PGM activity in maturing *USP::TPP* and WT embryos. Values are means ± SEM (n=25), * P ≤ 0.05 (Student’s t-test). (**B**) Relative abundance of PGM2 transcripts in *USP::TPP* and WT embryos over a period from 14 to 26 DAP. Values given as means ± SEM (n=10).

**Supplemental Figure 4.**
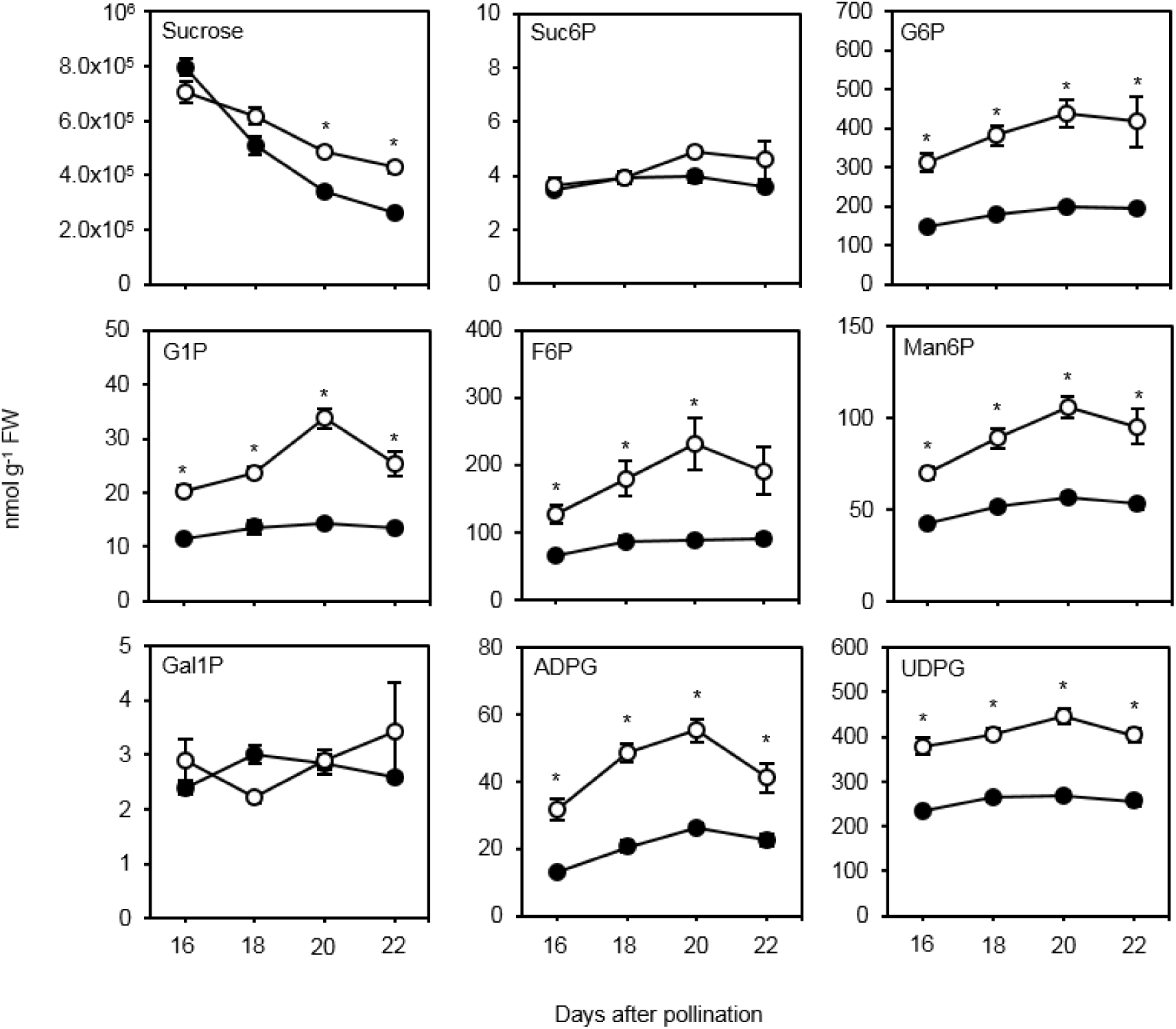
Heterologous expression of TPS in pea embryos affects sucrose and sugar phosphate concentrations. Soluble sugar levels of developing *USP::TPS* (solid circles) and WT (open circles) embryos at 16, 18, 20, and 22 DAP. Error bars, SEM (n=15); significance was calculated according to Student’s t-test: *P ≤ 0.001. ADPG, ADP-glucose; F6P, fructose 6-phosphate; G1P, glucose 1-phosphate; G6P, glucose 6-phosphate; Gal1P, galactose 1-phosphate; Man6P, mannose 6-phosphate; Suc6P, sucrose 6’-phosphate; UDPG, UDP-glucose.

**Supplemental Table 1.** TPP activity and T6P content in 16-day-old embryos of five homozygous *USP::TPP* and their corresponding WT lines.

| Plant line | TPP activity | T6P content |
| --- | --- | --- |
| | $\mu\text{mol g}^{-1} \text{FW} \cdot \text{min}^{-1}$ | $\text{nmol g}^{-1} \text{FW}$ |
| *USP::TPP* #1 | 0.41 ± 0.02 | 0.70 ± 0.04* |
| WT #1 | ND | 2.50 ± 0.17 |
| *USP::TPP* #2 | 0.54 ± 0.02 | 0.67 ± 0.04* |
| WT #2 | ND | 2.24 ± 0.17 |
| *USP::TPP* #3 | 0.43 ± 0.04 | 0.93 ± 0.07* |
| WT #3 | ND | 2.96 ± 0.11 |
| *USP::TPP* #5 | 0.49 ± 0.04 | 0.74 ± 0.04* |
| WT #5 | ND | 2.07 ± 0.12 |
| *USP::TPP* #6 | 0.54 ± 0.02 | 0.62 ± 0.01* |
| WT #6 | ND | 2.99 ± 0.19 |

Values given as means ± SEM ( $n=5$ ). \* $P \leq 0.001$ (Student's $t$ -test) ND, not detectable

**Supplemental Table 2.** Compositional analysis of mature seeds harvested from transgenic *USP::TPP* and corresponding WT plants.

|  | USP::TPP | WT |
| --- | --- | --- |
| Seed weight (mg) | 199 ± 5* | 319 ± 7 |
| Starch (mg g<sup>-1</sup> DW) | 348 ± 3* | 435 ± 4 |
| Starch (mg per seed) | 69.2 ± 2.0* | 139 ± 3 |
| Sucrose (μmol g<sup>-1</sup> DW) | 84.8 ± 6.7* | 64.8 ± 2.0 |
| Total N (%) | 4.6 ± 0.1* | 4.0 ± 0.07 |
| Total C (%) | 44.2 ± 0.04* | 44.5 ± 0.01 |
| C/N ratio | 9.7 ± 0.2* | 11.1 ± 0.3 |

Values given as means ± SEM (*n*=5). \**P* ≤ 0.05 (Student's *t*-test)

**Supplemental Table 3.** The T6P content of 16-day-old *USP::TPS* and the corresponding WT embryos.

|  | USP::TPP | WT |
| --- | --- | --- |
| Cell area ($\mu\text{m}^2$) | 7.93 ± 0.84* | 9.3 ± 0.68 |
| Cotyledon length (mm) | 5.23 ± 0.37* | 6.77 ± 0.22 |
| Cotyledon width (mm) | 2.83 ± 0.08* | 2.62 ± 0.05 |
| Cotyledon length/width ratio | 1.86 ± 0.16* | 2.59 ± 0.12 |

Values given as means ± SEM ( $n=5$ ). \* $P \leq 0.05$ (Student's $t$ -test)

**Supplemental Table 4.** The level of soluble sugars in 24 DAP embryos harvested from r, rb, and rug4 mutant plants.

|  | *r* | *R* | *rb* | *Rb* | *rug4* | *RUG4* |
| --- | --- | --- | --- | --- | --- | --- |
|  | *nmol g<sup>-1</sup> FW* |  |  |  |  |  |
| ADPG | 182 ± 17* | 35.2 ± 3.1 | 9.1 ± 0.4* | 15.5 ± 1.6 | 5.5 ± 0.7* | 23.6 ± 2.2 |
| UDPG | 193 ± 16* | 241 ± 27 | 204 ± 6* | 135 ± 18 | 25.6 ± 3.2* | 193 ± 12 |
| G6P | 1185 ± 44* | 320 ± 13 | 931 ± 27* | 220 ± 10 | 184 ± 3* | 253 ± 5 |
| G1P | 69.8 ± 2.3* | 23.7 ± 0.6 | 55.2 ± 1.4* | 18.2 ± 1.0 | 13.0 ± 0.4* | 19.7 ± 0.5 |
| F6P | 329 ± 16* | 116 ± 8 | 290 ± 6* | 78.2 ± 7.0 | 59.0 ± 2.5* | 80.5 ± 5.6 |

Values given as means ± SEM (*n*=5). \**P* ≤ 0.05 (Student's *t*-test)
ADPG, ADP-glucose; F6P, fructose 6-phosphate; G1P, glucose 1-phosphate; G6P, glucose 6-phosphate; UDPG, UDP-glucose

**Supplemental Table 5.** The extent of AGP monomerization in 18-, 22-, and 26-day-old DAP embryos, harvested from plants harboring the *USP::TPP* transgene.

| Plant line | AGP monomerisation (%) |  |  |
| --- | --- | --- | --- |
|  | 18 DAP | 22 DAP | 26 DAP |
| *USP::TPP* #1 | 45 ± 0 | 25 ± 1 | 41 ± 1 |
| WT #1 | 41 ± 1 | 29 ± 3 | 42 ± 1 |
| *USP::TPP* #2 | 58 ± 6 | 31 ± 2 | 34 ± 2 |
| WT #2 | 49 ± 1 | 27 ± 1 | 32 ± 3 |
| *USP::TPP* #3 | 41 ± 1 | 13 ± 2 | 31 ± 2 |
| WT #3 | 39 ± 0 | 20 ± 4 | 36 ± 5 |
| *USP::TPP* #5 | 58 ± 0 | 16 ± 3* | 38 ± 3 |
| WT #5 | 53 ± 3 | 30 ± 4 | 40 ± 2 |
| *USP::TPP* #6 | 54 ± 3 | 13 ± 3* | 27 ± 3 |
| WT #6 | 44 ± 2 | 30 ± 4 | 32 ± 3 |

Values given as means ± SEM ( $n=5$ ). \* $P \leq 0.05$ (Student's $t$ -test)

**Supplemental Table 6.** TPP activities in 26-day-old embryos formed by hybrids between *USP::TPP* and *USP::TAR2* plants.

| Plant line | TPP activity |
| --- | --- |
| | $\mu\text{mol g}^{-1} \text{FW} \cdot \text{min}^{-1}$ |
| Empty vector | ND |
| *USP::TPP* #2 | 0.15 ± 0.03 |
| *USP::TPP* #3 | 0.17 ± 0.03 |
| *USP::TAR2* #3 | ND |
| *USP::TAR2* #5 | ND |
| *USP::TPP* #2 / *USP::TAR2* #3 | 0.17 ± 0.02 |
| *USP::TPP* #3 / *USP::TAR2* #3 | 0.14 ± 0.02 |
| *USP::TPP* #3 / *USP::TAR2* #5 | 0.14 ± 0.03 |

Values given as means ± SEM ( $n=3$ ). ND, not detectable

**Supplemental Table 7.**
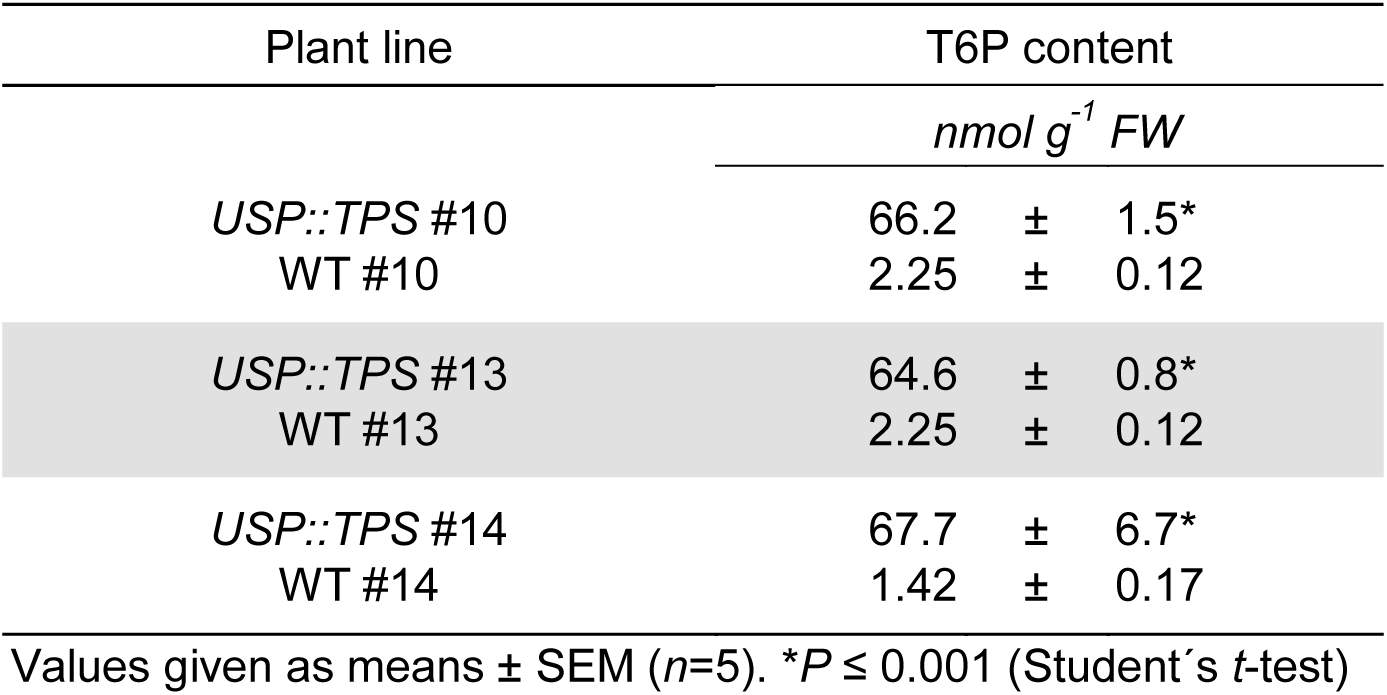
The T6P content of 16-day-old *USP::TPS* and the corresponding WT embryos.

| Plant line | T6P content |  |  |
| --- | --- | --- | --- |
|  | *nmol g<sup>-1</sup> FW* |  |  |
| *USP::TPS* #10 | 66.2 | ± | 1.5* |
| WT #10 | 2.25 | ± | 0.12 |
| *USP::TPS* #13 | 64.6 | ± | 0.8* |
| WT #13 | 2.25 | ± | 0.12 |
| *USP::TPS* #14 | 67.7 | ± | 6.7* |
| WT #14 | 1.42 | ± | 0.17 |

Values given as means ± SEM (*n*=5). \**P* ≤ 0.001 (Student's *t*-test)

**Supplemental Table 8.** Compositional analysis of mature seeds harvested from transgenic *USP::TPS* and corresponding WT plants.

|  | USP::TPS | WT |
| --- | --- | --- |
| Seed weight (mg) | 325 ± 4 | 321 ± 8 |
| Starch (mg g<sup>-1</sup> DW) | 437 ± 8 | 460 ± 6 |
| Sucrose (μmol g<sup>-1</sup> DW) | 45.5 ± 2.9* | 76.2 ± 5.1 |
| Total N (%) | 3.7 ± 0.1 | 3.4 ± 0.1 |
| Total C (%) | 44.5 ± 0.0* | 44.2 ± 0.0 |
| C/N ratio | 12.0 ± 0.3 | 13.2 ± 0.4 |

Values given as means ± SEM (*n*=5). \**P* ≤ 0.001 (Student's *t*-test)

**Supplemental Table 9.**
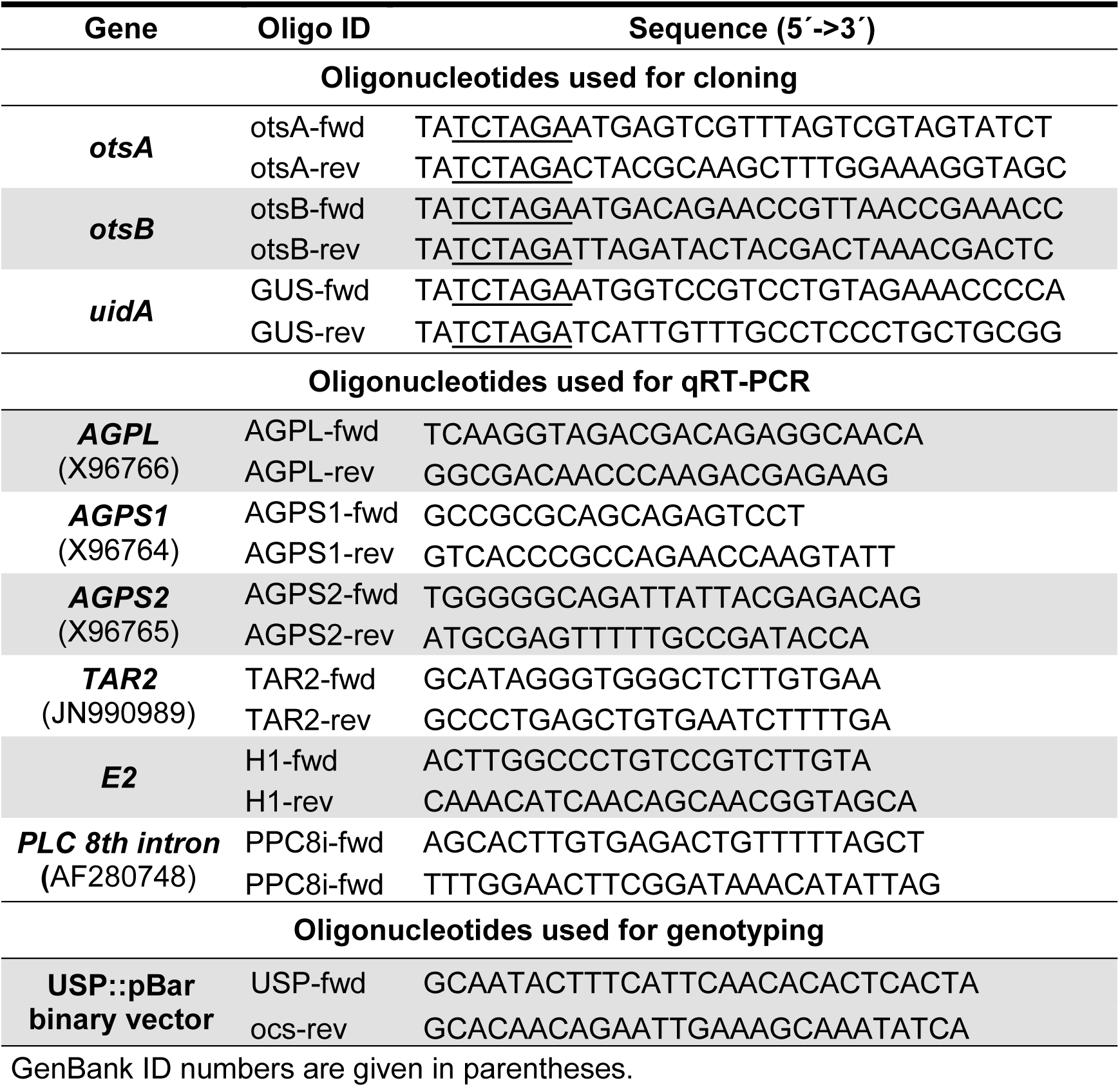
Oligonucleotides sequences employed.

